# What shapes template matching performance in cryogenic electron tomography *in situ*

**DOI:** 10.1101/2023.09.06.556487

**Authors:** Valentin J. Maurer, Marc Siggel, Jan Kosinski

**Author notes:** Contributed equally (Valentin J. Maurer, Marc Siggel). Electronic mail (Valentin J. Maurer, Jan Kosinski).

## Abstract

Detecting specific biological macromolecules in cryogenic electron tomography (cryo-ET) data is frequently approached by applying cross-correlation-based 3D template matching. To reduce computational cost and noise, high binning is used to aggregate voxels before template matching. This remains a prevalent practice in both practical applications and method development. Here, we systematically evaluate the relation between template size, shape, and angular sampling to identify ribosomes in a ground truth annotated dataset. We show that at the commonly used binning, a detailed subtomogram average, a sphere, and the heart emoji 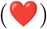 results in near-identical performance. Our findings indicate that with current template-matching practices, macromolecules can only be detected with high precision if their shape and size are sufficiently different from the background. Using theoretical considerations we rationalize our experimental results and discuss why at high binning, primarily low-frequency information remains and that template matching fails to be accurate because similarly shaped and sized macromolecules have similar low-frequency spectrums. We discuss these challenges and propose potential enhancements for future template-matching methodologies.

## I. NTRODUCTION

Cellular cryogenic electron tomography (cryo-ET) has emerged as a key method to unravel the structural and spatial complexity of the cell. The 3D volume of a cellular region, called a tomogram, is reconstructed from 2D projection images acquired on a transmission electron microscope in many different orientations.^1,2^ Macromolecular complexes can be identified in the tomogram, their spatial arrangement can be analyzed in the native environment, and their structure can potentially be determined to near-atomic resolution by subtomogram averaging.^3–8^ However, identifying individual macromolecules in tomograms is challenging due to the missing wedge, which results from limitations on possible tilt angles of the specimen, low signal-to-noise ratio due to low electron doses used during acquisitions, and the crowded cellular environment of the cell. Because of these complications, segmentation and subsequent analysis of tomograms is still a difficult task and a major bottleneck for fully automated high-throughput analysis of large cryoET datasets.^1,6,9,10^

A commonly used approach to identify macromolecules within tomograms is so-called template matching, where a reference density of a macromolecule is used to localize the corresponding candidate positions within the tomogram.^11,12^ To date, various packages have been developed to perform template matching such as PyTom^13^, STOPGAP^14,15^, EMAN2^16^, DYNAMO^17^, and pyTME^18^. All of these packages use cross-correlation-based scoring metrics to identify macromolecules in tomograms (see III D). Templates used for template matching range from simple shapes such as spheres, cylinders, and rectangles, which have been previously used to detect various cellular structures^19–22^, to detailed maps obtained in experiments or generated from atomic structures.^11,23,24^ A common use case is the ribosome, which is abundantly found in tomograms and can often be identified by eye due to its size. However, even for a particle as large as the ribosome, template matching has suboptimal precision.^10,25^ For smaller or less abundant macromolecules, the method often fares even worse and requires manual curation. These points raise the question of what the requirements and limitations are to reliably use template matching for macromolecules in *in situ* cryo-ET.

Although tomograms are usually collected at 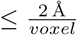, they are typically binned 4-8 times to a coarse voxelsize in order to improve computational efficiency and signalto-noise ratio.^7,8,10,11,15,19,20,26–28^ However, binning removes high-frequency information from the tomogram and makes it difficult to distinguish macromolecules if the differences in low-frequency components are not sufficiently large.^12^ Therefore, template matching under such settings is prone to have low precision, and manual curation or other means of refinement are required to improve the results. The issue of low precision has been hinted at previously, and one suggested solution is template matching in 2D.^29,30^ However, to our knowledge, there has been no published study that systematically explored how the choice of an exact template, its size, and the degree of angular sampling affects *in situ* template matching results.

Here, we investigate the pitfalls of 3D template matching with commonly used four times binned tomograms and rationalize the observed issues. We assess the precision of detecting ribosomes in a previously annotated tomogram^10^, using a detailed subtomgram average of a ribosome, a sphere, an emoji 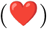, and a structure of hemagglutinin, all of varying radii as templates. We find that at this binning, the size and approximate shape are the major determinants of precision, and exact template choice or angular sampling has little impact on the template matching results. We then rationalize these observations theoretically by inspecting the Fourier transforms of simple geometric shapes and show that, when low-frequency components dominate in the tomogram, similarly sized and shaped templates result in nearly identical template matching precision. Finally, we discuss the implications of these results for the development and benchmarking of template-matching algorithms as well as requirements for data processing. Further, we aim that our analysis will guide optimal experimental design in practical applications and the development of new templatematching methods.

## II. METHODS

All template matching experiments were performed using PyTom^13^ (version 1.0), and cross-validated using pyTME^18^ (version 0.1.7), on an annotated tomogram (EMPIAR-10988, TS_037, ref. 31) reported by de Teresa-Trueba *et al*. ^10^. For this sample tomogram, 1.646 ribosomal and 22 fatty acid synthase (FAS) positions were previously identified using PyTom^13^ and subsequently manually refined by an expert user. The template matching was performed for four different template classes that were provided in varying sizes, as shown in Fig. 1. First, a previously reported map of the *S. cerevisiae* 80S ribosome (EMD-3228; ref. 32) was used as a baseline reference (Fig. 1A) with an approximate radius of ten voxels.^10^ The structure was scaled to match radii between 1 and 19 voxels using linear spline interpolation. Second, a sphere with varying radius *r* 1 ≤ *r <* 20 and homogenous density (Fig. 1B) was used. The third template was the emoji 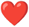 from the Apple Color Emoji font. The 2D bytemap was converted to a volume with homogenous density by axial symmetrization sampling 360 equidistant angles and subsequently blurred using a Gaussian filter (scipy.ndimage.gaussian_filter, version 1.11.1) with *σ* = 1 (Fig. 1C). The heart emoji was scaled from the initial 160 × 160 bytemap to appropriate radii using linear spline interpolation. As an additional control template with a clearly distinct shape and size, a structural model of the hemagglutinin trimer was used.

**FIG. 1.**
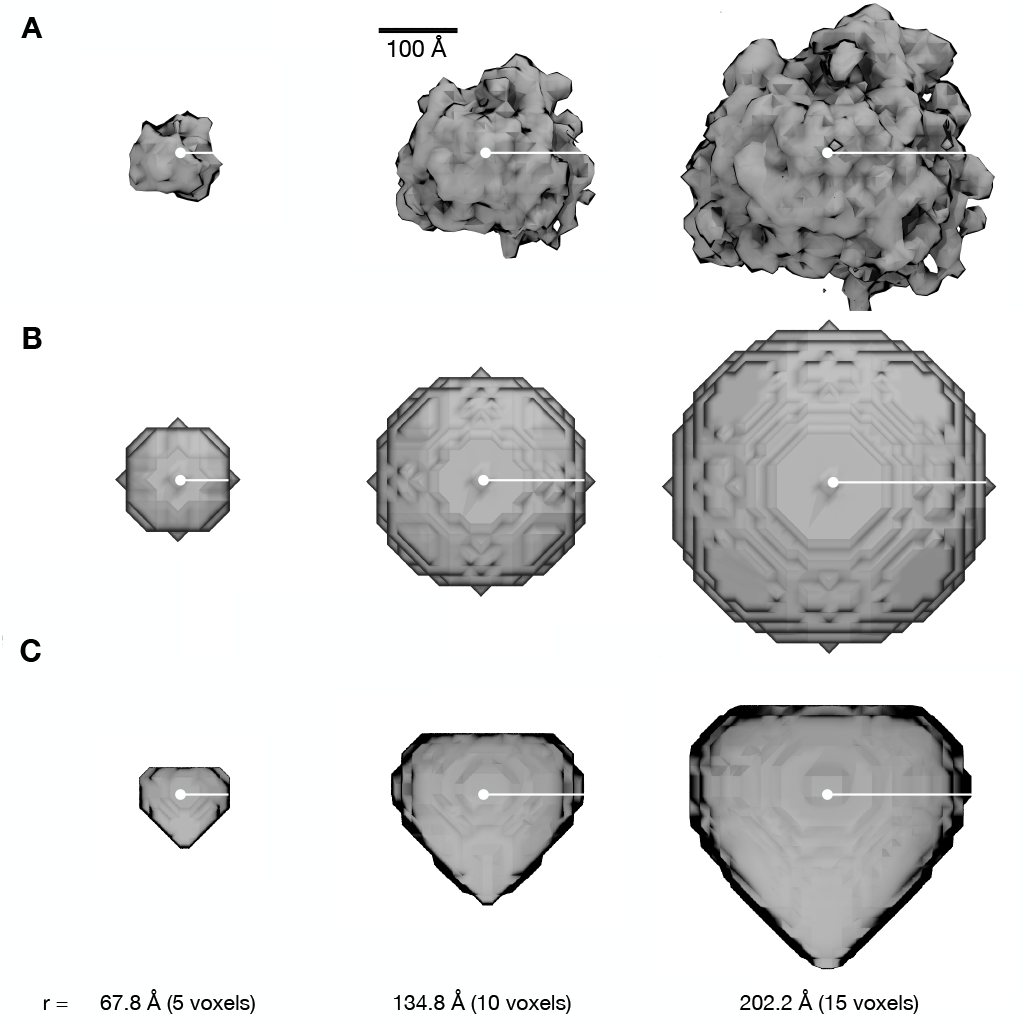
Template classes used to match the ribosome in the previously annotated tomogram from de Teresa-Trueba *et al*. ^10^. Different shapes with different radii sampled at a 13,48 Å voxel size, to match the voxel size of the tomogram, were used as templates for template matching with PyTom. Specifically, the map of the 80S ribosome (**A**, EMD-3228), a sphere (**B**), and a rotationally symmetrized emoji, the heart emoji, (**C**) were used. The used radii range from 1 to 19 voxels in one voxel increments. Only three representative radii are shown.

The atomic structure was obtained by modeling with AlphaFold2^33^ multimer using the A/Hong Kong/1/1968 H3N2 strain. The default parameters were used with an exception for the number of refinement cycles being increased to 6. The best model was chosen based on the lowest overall prediction alignment error. The atomic structure was discretized on a grid with the sampling rate of the tomogram (Fig. S1A). To simulate different radii, the grid was resampled to a sampling rate computed as 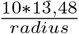, i.e. a radius of 11 corresponds to 1,1 times the sampling rate of the tomogram. The structure was not discretized on grids of varying sampling rates directly, to avoid the introduction of additional detail for higher radii. All templates were placed in the center of a cubical volume with edge length 51 and voxel size 13,48 Å, which corresponds to the voxel size of the used tomogram. As a mask, a sphere whose radius was two voxels larger than the template radius was used, which is in good concordance with the 337 Å diameter mask used by de Teresa-Trueba *et al*. ^10^. Wedge correction was performed using 40° angles, which corresponds to the -50° to 50° tilt acquisition angles.^10^ PyTom was used to sample translations exhaustively at voxel size resolution. For the spheres, 90° (two angles), and for the other templates, 25.25° (980 angles), 19.95° (1.944 angles), and 11° (15.192 angles) were sampled, which corresponds to the angle lists angles_90_2, angles_25.25_980, angles_19.95.25_1944 and angles_11_15192, that are shipped with PyTom.^13^ The rotational sampling rate of 11° is in excess of what is typically used for template matching in previous work.^10,13,34^ PyTom was run assuming a spherical mask, used a bandpass filter with a low and high-frequency cutoff of 3 and 15, split the tomogram into 3 parts along each axis, and performed no further binning. The results were cross-validated using pyTME, which can perform template matching using a similar formulation of the crosscorrelation score.^18^ pyTME was run on the same data, with the difference that no bandpass filter and no wedge mask were applied prior to template matching. 4.000 peaks were called on the scores obtained from PyTom and pyTME using skimage.feature.peak_local_max (version 0.21.0) with a minimal allowed Euclidean distance separating peaks of 10 and a 15 voxel exclusion volume around the boundaries of the tomogram. Subsequently, peaks were ordered by their score in decreasing order. The precision and recall were analyzed at decreasing score thresholds up to 4.000 top-scoring peaks.

## III. RESULTS AND DISCUSSION

### A. Shape and size are the major determinants for template matching precision

To assess how sensitive template matching is *in situ* to a template’s specific shape and size, we used PyTom^13^ to perform template matching with four different templates on a four-times binned tomogram with a voxel size of 13,48 Å reported in de Teresa-Trueba *et al*. ^10^. This is comparable to the voxel sizes typically used in many previously published template matching studies^7,8,10,11,15,19,20,26,27^. We also used pyTME^18^ to independently cross-validate these results. The tomogram contained 1.646 ribosomes and 22 FAS particles, which were annotated by the authors using template matching and manual curation. We consider their annotation as robust ground truth. As per de Teresa-Trueba *et al*. ^10^, we used an 80S ribosome^32^ map as the initial template and scaled its radius to see how size affects template matching performance (Fig. 1A). We also tested spheres and an irregularly shaped heart emoji at various radii (Fig. 1B,C) to check whether the exact properties of the template are relevant at this binning to achieve high precision in template matching and also compared a variety of angular sampling rates. Spheres have already been successfully used in practice for identifying RuBisCO^19^ and other basic shapes such as cylinders for nucleosomes^20^ or the proteasome^21^ and rectangles for membranes.^22^ However, a side-by-side comparison has been lacking to date. Therefore, we compared the template matching results for the different templates and scaled radii and angular samplings based on precision 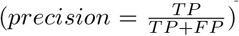 and recall 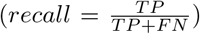, where *TP, FP*, and *FN* correspond to the number of true positives, false positives, and false negatives, respectively (Fig. 2). The picked particles were chosen from a sorted list of all scores in descending order.

**FIG. 2.**
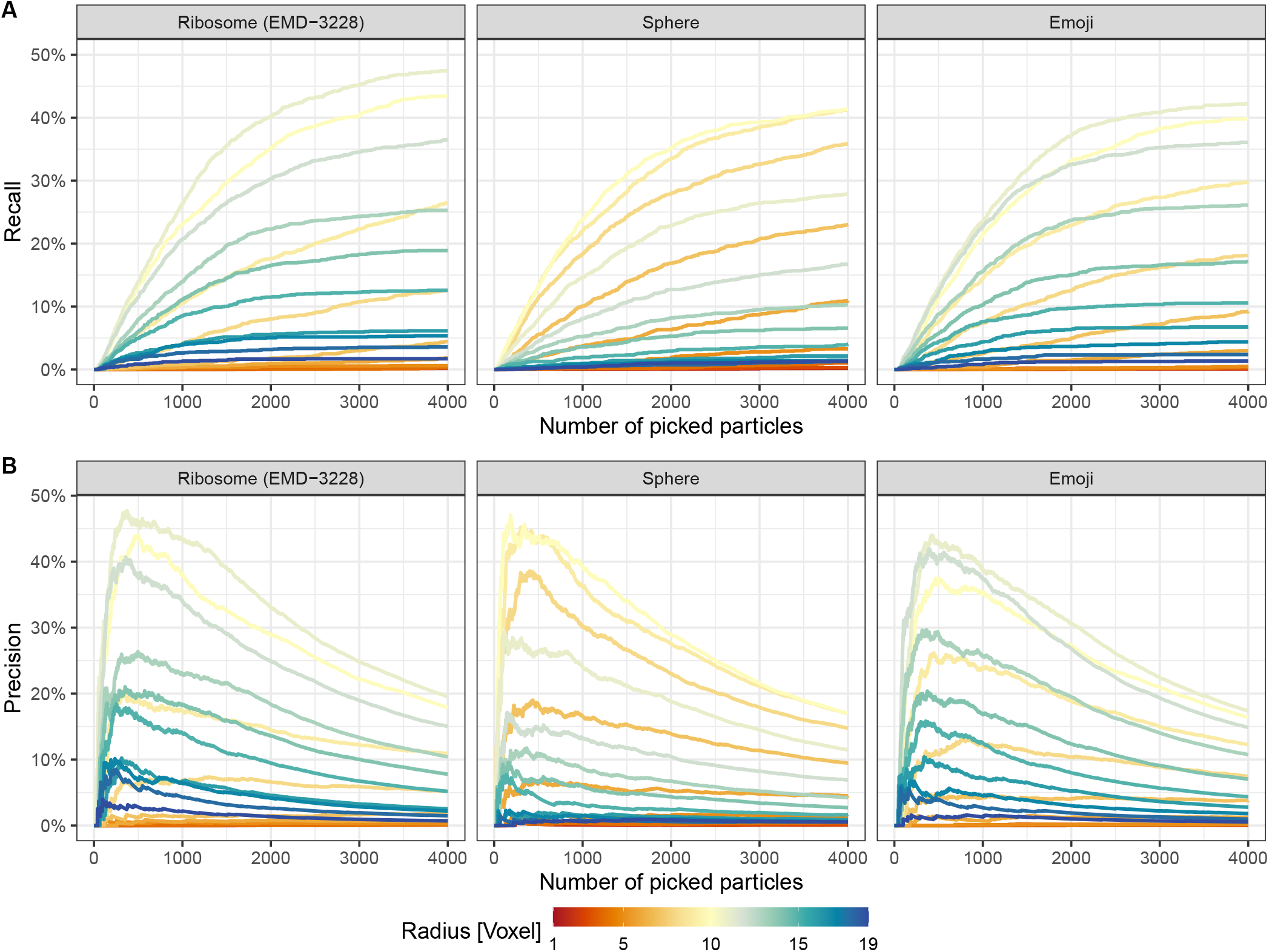
Template matching performance using three distinct templates (see Fig. 1). **A** Ribosome picking recall by number of picked particles. We used linear sum assignment to achieve an optimal one-to-one mapping between ground truth and picked particles. A particle is considered correctly picked if it is within a 20-voxel distance from its assigned ground truth particle. Consequently, all particles without assignment to ground truth particles were considered false positives. All 4.000 picked particles were considered in the following figures. **B** Same as in **A** but showing the precision instead of recall.

First, we compared the recall across the different templates and with different radii calculated with respect to the number of picked particles (Fig 2A). Overall, the performance of up to 4.000 top-scoring picks across templates was comparable, with a maximal recall of around 40-50%. At the voxel size of 13,48 Å used here, a realistic *S. cerevisiae* ribosome map (EMD-3228; ref. 32) performs no better than a sphere or an emoji of similar size on the same dataset. Furthermore, given the shape of the curves, it appears unlikely that the remaining ribosomes could be recovered at reasonable precision. Given that no template recapitulated the ground truth particle set beyond a recall of 50%, it becomes questionable whether using improved experimental or predicted structures as templates will be sufficient to identify small proteins or low abundance proteins in cryo-ET data at this binning. This is further substantiated by ribosomes requiring manual curation^10^. Likely, other high-scoring picks correspond to other particles or features of similar shape and size.

Similarly, the precision for the different templates peaked at around ≈ 750 picked particles independent of the template choice (Fig. 2B). Picking more than 750 particles improved the overall recall but led to a disproportionate identification of false positives, thus reducing the overall precision. We observed this behavior consistently for all templates, and there was little differentiation between the correct ribosome template and the sphere or emoji template. The precision values observed here are in line with the 19 % reported by de Teresa-Trueba *et al*. ^10^ for all ten defocus tomograms.

To further confirm this finding, we also ran control experiments using an Influenza A virus hemagglutinin (HA) template, which has a markedly different shape than a ribosome (Fig. S1A). HA is a trimer with a total molecular weight of 180 kDa with an approximately cylindrical shape with a length of ≈17 nm and a width of ≈6 nm. We scaled the radius analogous to the previous structures and calculated the precision with respect to the ground truth data. The precision was near 0% for sizes up to 10 voxels, and only for larger radii the precision increased, as the structure further approaches the shape and size of the ribosome (Fig. S1B). This further underscores the observation that at this level of binning, template matching is less dependent on the structure and overall focuses on shape and size.

When comparing the different template radii, we observed that the radius, not the chosen template had the largest effect on the overall precision (Fig. 2B and 3). The template matching precision at 4.000 picks was maximal at around 10-11 voxels. Templates smaller than a radius of ten voxels performed considerably worse. This is likely due to the presence of noise or additional macromolecules that are smaller than the ribosomes but are composed of comparable density. This is in line with the fact that many studies perform template matching with large macromolecules including, but not limited to, ribosomes^10,13,26,34,35^, proteasomes^11,21^, thermosomes^11^, RuBisCO.^19^ We also note that these results were independent of the software used, as pyTom and pyTME resulted in near-identical precision (Fig. 3).

**FIG. 3.**
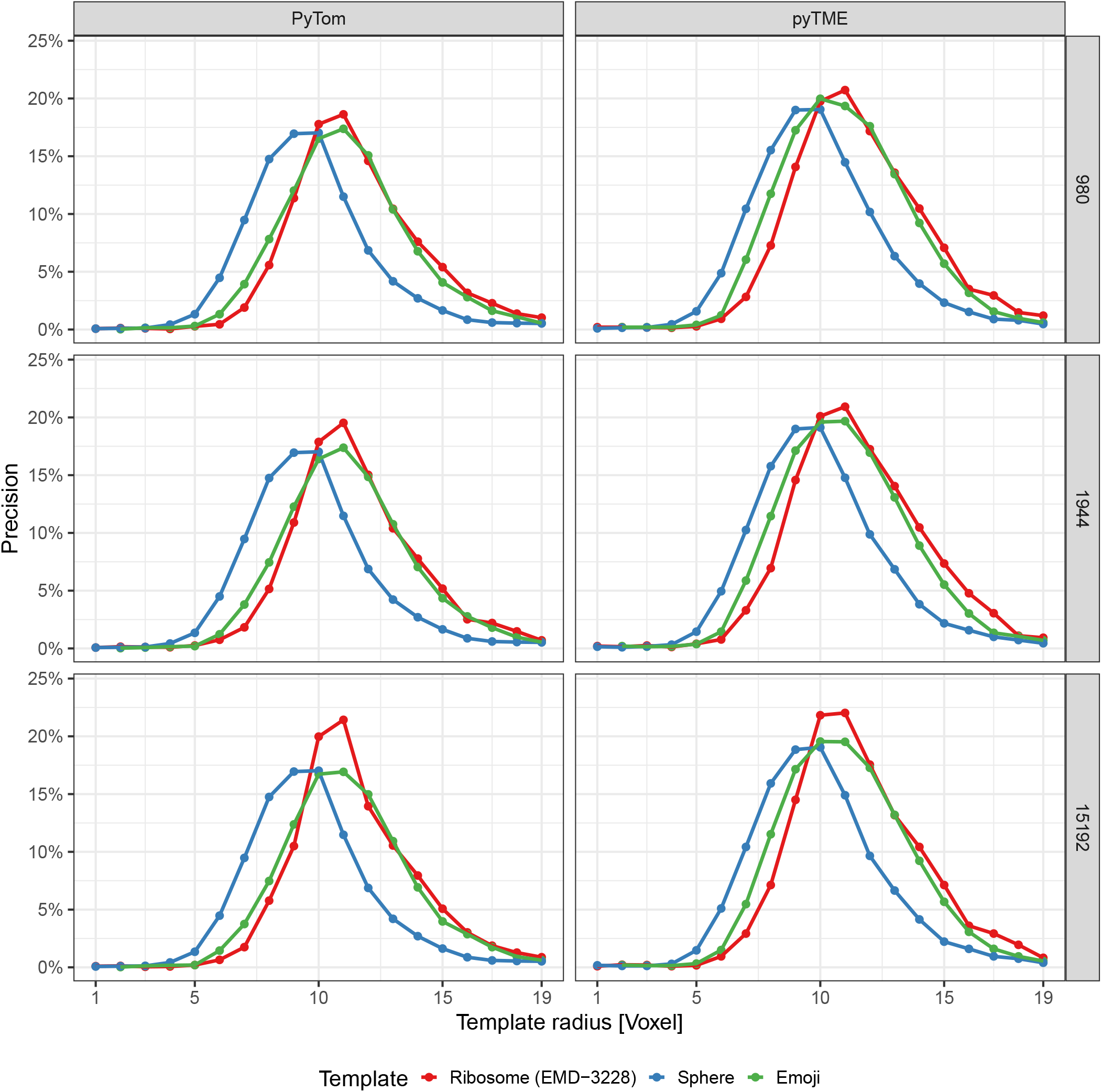
Proportion of true positives out of all picked particles (Precision) by radius of templates. Shown in the left column are template-matching results obtained with the software PyTom on the left and with pyTME on the right. In each row, the sampled number of angles is shown, namely 980, 1,944, and 15,192. 4.000 picks are considered when determining the precision.

### B. Angular sampling does not improve precision

We also tested the effect of varying angular sampling on the result to ensure that undersampling did not affect the results (Fig. 3). Higher angular sampling with 15.192 angles, compared to the 1.944 angles used above, did not significantly change the differentiation between the shapes and only increased the precision by approximately 3%. This indicates that while for purified, *in vitro* samples^26^ higher angular sampling improves results at a 13 Å voxel size, *in situ* samples do not benefit from higher angular sampling to the same extent. In this case, an increase in precision by 5% does not warrant the use of approximately 15 times more computational resources. This is also to be expected since the cross-correlation score does not scale exponentially, and subtle increases in the score do not necessarily increase differentiation from similar-sized and shaped objects in the *in situ* sample.

We also cross-validated these results with software package pyTME^18^ to ensure that software-specific implementation details did not affect this result. With both packages, the results were near-identical across all sampled conditions.

Based on these findings, we suggest initially filtering candidate positions with low angular sampling, using potentially even a simple shape-based template of appropriate size, and sampling the same positions at a lower binning or removing false positive hits through classification with other tools such as RELION^36^. Such workflows have been previously proposed in packages such as nextPyP^37^, tomoBEAR^38^, or Dynamo.^39^ Specifically in this case, using a spherical template is computationally most efficient because it is rotationally invariant and thus yields the best time-to-solution as it can be run without any angular sampling.

### C. Ribosome and Fatty Acid Synthase are not discernible with conventional template matching

Finally, we assessed whether using the ribosome, sphere, and emoji templates at different scales, we could pick the FAS protein complex, whose shape differs substantially from the ribosome’s shape but has a similar size. Although the number of annotated FAS in the particular tomogram is only 22, FAS particles are among the 4.000 highest-scoring picks across templates and scales (Fig. 4). Although the low number of annotated FAS impedes quantitative claims, the trends are clear. For a sphere, as many as 40% of FAS are recovered, and even for the ribosome as a template, more than 45% of annotated FAS instances are recovered. This finding further highlights that molecular details play a minor role in template matching at our voxel size of 13,48 Å. In our case FAS and ribosomes are similarly sized, resulting in fairly similar scores and, thus, poor differentiation between the two. Generally, this indicates that low-abundance proteins cannot be practically identified with sufficient precision if many other macromolecules of similar size exist.

**FIG. 4.**
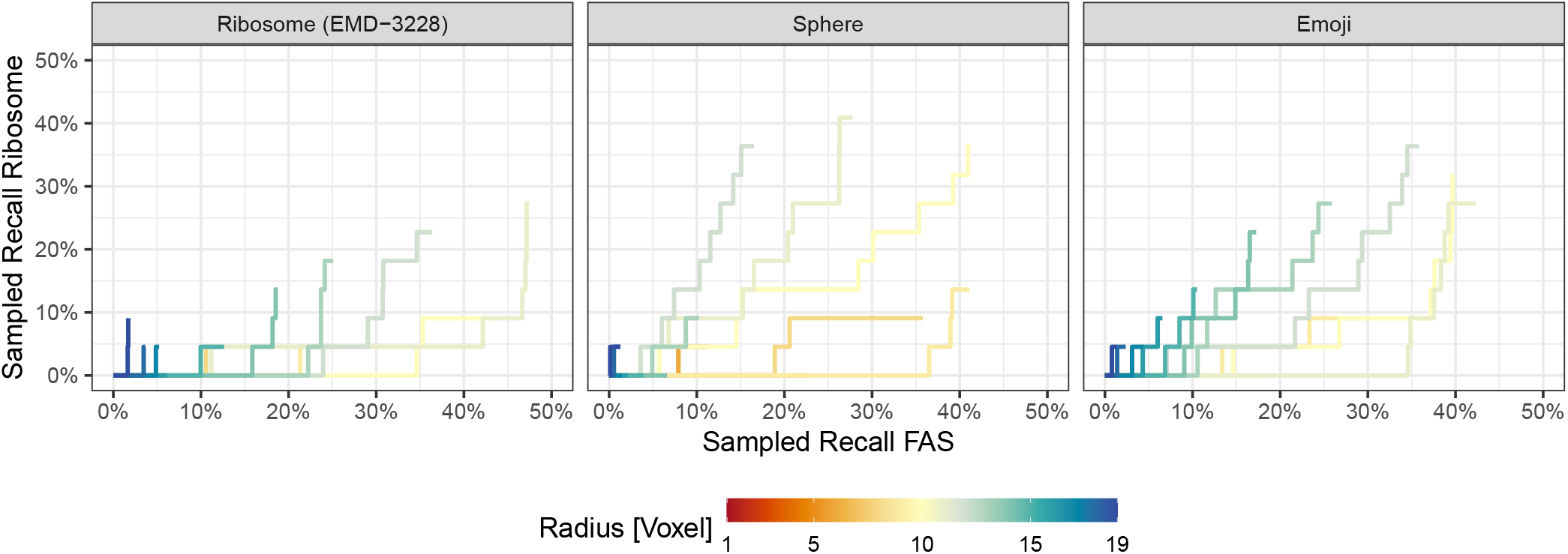
Template matching performance on the FAS complex using three distinct templates. Picked particles were one-to-one assigned to the union of ground truth FAS and ribosome coordinates using linear-sum assignment. Each particle is assigned to no more than one class and considered to correctly pick that class if it is within a 20-voxel distance from its assigned ground truth particle.

### D. Theory

We now aim to rationalize our empirical observations by examining the analytical form of the Fourier transforms of several geometric shapes and discussing them in the context of cross-correlation calculation. Based on our assessment, we conclude that template matching on the typically used 4-8 times binning is primarily driven by shape and size and list the associated implications.

Most template matching programs, including PyTom^13^ and pyTME^18^, use the cross-correlation theorem to determine the similarity between a target *f* and a template *g* at a given translation *n*:

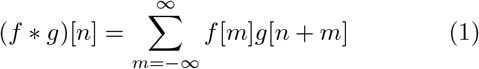

Where ∗ is the convolution operator. Cross-correlation is the sum of the element-wise product of the template and the target, subject to implementation-specific normalization procedures. In practice, this procedure is repeated for a set of rotations of the template.

The computational complexity of the cross-correlation operation on two identical cubes with edge length N is *𝒪* (*N* ^6^), but in practice, template matching tools reduce the complexity to *𝒪* (*N* ^3^ log(*N* ^3^)). This is achieved by expressing the cross-correlation in the spatial domain as multiplication in the Fourier domain through the crosscorrelation theorem:

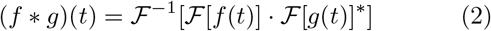

Where *ℱ* and *ℱ*^−1^ denote the forward and inverse Fourier transform and ^∗^ the complex conjugate. To build some intuition on how this impacts template matching, let’s consider the case *g*(*t*) = *f* (*t* − *n*), where *g* differs from *f* only by a translation *n*:

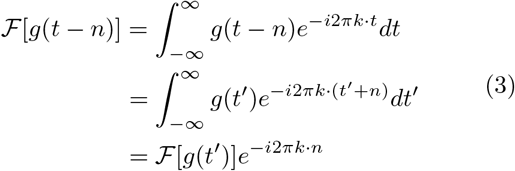

where *k* is the wave number in Fourier, and *t* is the position vector in the real domain. From this, it becomes apparent that a shift in the spatial domain corresponds to a frequency-dependent phase shift in the Fourier domain. Furthermore, we note that since:

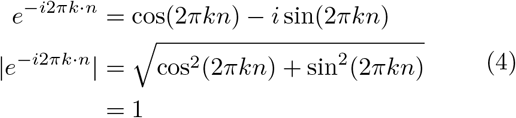

the magnitude of the Fourier transform is independent of the phase shift. Therefore, we can determine the crosscorrelation in the real domain by inverse Fourier transform of the element-wise product of amplitudes and the sum of phases of *ℱ* [*f* ] and *ℱ* [*g*]^∗^. The magnitude of the cross-correlation score depends on the element-wise product of amplitudes, while the sum of phases contains the mapping between translation and score, which is maximal at translation *n*.

Above, we considered the ideal case where the template is a shifted version of the target. In practice, this rarely holds, and the template rather approximates the amplitude and shifted phase spectrum of the target sufficiently well. Therefore, previous research has seen the use of geometric shapes for template matching, such as spheres for localization of ribosomes or RuBisCO^19^, cylinders for nucleosomes^20^ or proteasomes^21^, and rectangles for membranes.^22^ Intuitively, geometric shapes can be used for template matching if they approximate the structure of interest sufficiently well in the given data. However, why that is the case has not been shown explicitly. We aim to do so in the following and start by deriving the Fourier transforms of the aforementioned geometric shapes.

A sphere of radius *R* centered around the origin can be defined in real space as:

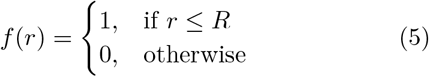

here *r* represents the magnitude of the position vector, i.e., the Euclidean distance from the origin. All points with a distance less than or equal to *R* are occupied by the sphere. Since the sphere is a real symmetric function, its Fourier transform is also real and follows as^40^:

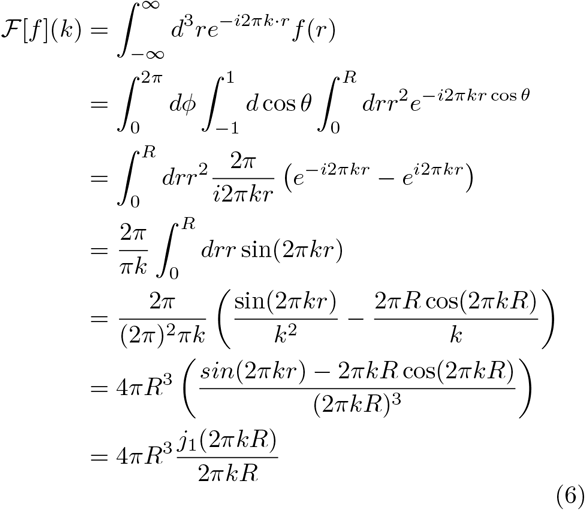

where *j*_1_(*x*) is the spherical Bessel function of first kind and order defined as:

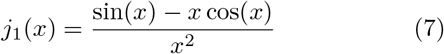

A one-dimensional rectangle, i.e., a box-function can be defined in real space as:

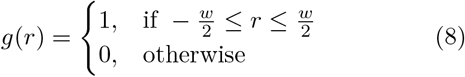

where *w* is the width of the box function. The Fourier transform of the one-dimensional box function g(r) is:

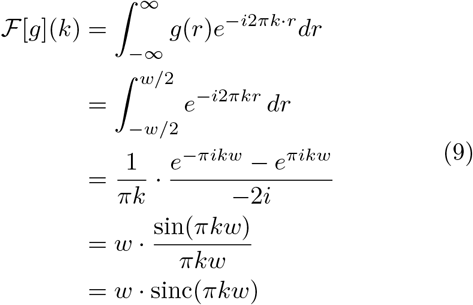

The definition of the box-function Fourier transform can be used to synthesize the Fourier transform of threedimensional rectangles with width a, b, and c as:

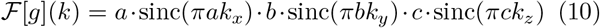

where *k*_*x*_, *k*_*y*_, and *k*_*z*_ are the wave numbers corresponding to the spatial dimensions x, y, and z, respectively.

The cylinder is essentially a combination of a circle and a box function and can be defined as follows:

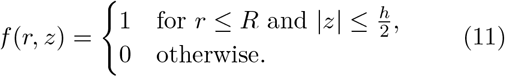

where *R* is the radius of the circle and *h* the width of the box-function. We can make use of the cylindrical symmetry, and the separability of the Fourier transform to derive the closed form of the cylinder Fourier transform as follows:

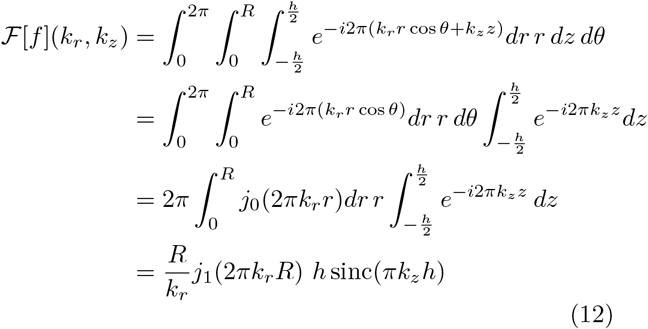

The Fourier transforms of the sphere, rectangle, or cylinder either contain a Bessel or a sinc function or a combination thereof. Therefore, these geometric shapes concentrate most of their Fourier energy in low-frequency components and dampen with a shape-specific rate toward higher frequencies. Böhm *et al*. ^12^ has already hinted at the fact that low frequencies are essential for particle identification and discussed the detection limits related to high binning.

To use geometric shapes in template matching, the macromolecule of interest within tomograms would also need to concentrate the majority of Fourier space energy in low-frequency components in a similar manner to yield a high cross-correlation score. Since low-frequency components generally recapitulate the shape and size of the analyzed object in real space, macromolecules have been template-matched by geometric shapes with similar sizes.^19–22^ The voxel size of the tomogram used in this study was 13,48 Å following common practices in the field.^7,8,10,11,15,19,20,26–28^ Therefore, no features smaller than ≈ 27 Å can be represented without artifacts according to the Shannon-Nyquist sampling theorem.^41^ 27 Å is in excess of most detailed structural features in a macromolecule, and consequently, the majority of Fourier space energy is also concentrated in low-frequency components, analogous to discussed the geometric shapes.

We computed the radially averaged Fourier power spectrum for three templates at a radius of ten voxels and compared them to the theoretical curve of a sphere (Fig. 5). All templates concentrate their Fourier domain energy in low-frequency components. This matches the theoretical assumption that template matching using geometric shapes is possible if the majority of the Fourier space energy is concentrated similarly. Accordingly, we see little differentiation in the total precision achieved for varying templates with the same radius and most variation between the same template and varying radii.

**FIG. 5.**
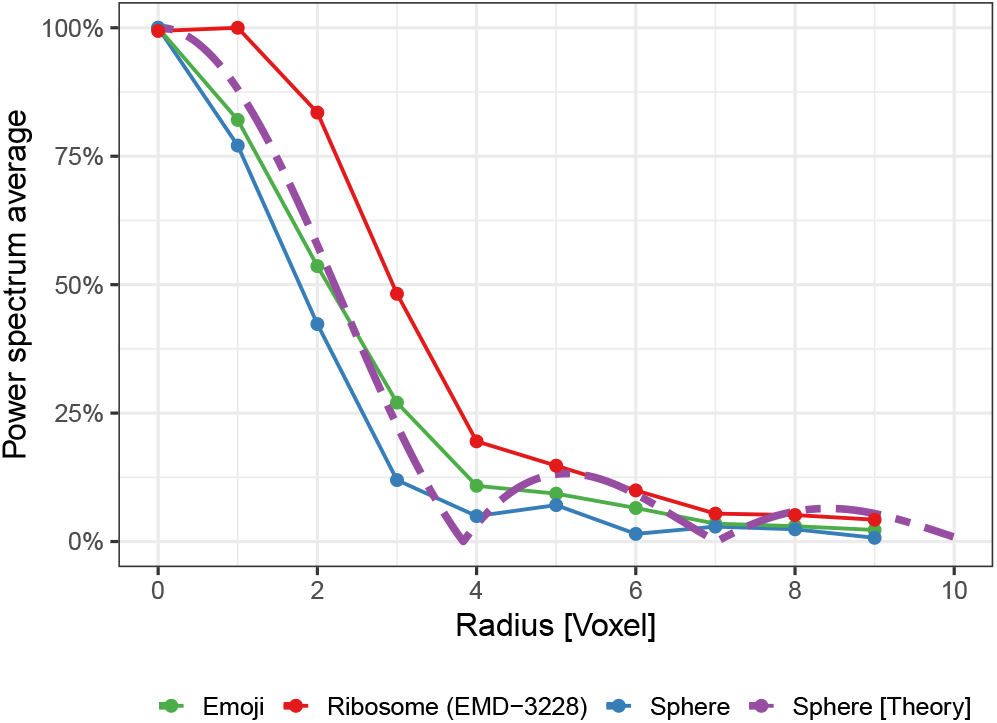
Fourier power spectrum averages by radius and template. “Radius” was computed as Euclidean distance from the zero-frequency component and rounded to the nearest integer. Shown are templates used for template matching at a radius of ten (Fig. 1). “Sphere [Theory]” refers to the theoretical derivations made in the theory section with *R* = 10.^40^ Power spectrum averages were linearly scaled to the interval [0, 1] to facilitate curve shape comparison.

These theoretical considerations have three important implications for template matching: (i) Since template matching at this binning is primarily about matching object size, macromolecules of similar size to the macromolecule of interest will be identified as false positives; (ii) Small macromolecules would be mainly represented through high-frequencies, which overlaps with noise in the data. This relation makes template matching small macromolecules at this binning near-impossible; (iii) More accurate templates are unlikely to improve template matching performance because high-resolution information cannot be accurately represented at the typically used 4-8 times binning.

### IV. CONCLUSIONS AND OUTLOOK

In this article, we explored the effect of shape, size, and angular sampling on the precision of matching ribosomes in an annotated *S. cerevisiae* tomogram at the commonly used 4-times binning.^7,8,1^ We showed that using a ribosome subtomogram average, a sphere, and the heart emoji as a template resulted in near-identical performance in our benchmark dataset, highlighting the shortcomings and limitations of using highly binned tomograms for template matching. We show, based on theoretical arguments, that because highly binned tomograms primarily consist of low-frequency information, geometric shapes such as spheres, cylinders, or rectangles of appropriate size can be used to identify macromolecules equally well as detailed structural templates. Therefore, cross-correlation scores are primarily driven by shape and size of the template, rather than internal structure, as seen in our practical experiments. This has important implications when moving to more complex datasets or smaller target structures in the future. At high binning, macromolecules of similar size will often be identified as false positives over the macromolecule of interest, regardless of how detailed the template is. Importantly, this issue will be more pronounced for small molecules where the high frequencies will overlap with noise, and template quality will not improve the performance either.

Based on these considerations, we suggest the following moving forward. First, cross-correlation-based scoring methods appear to be a suboptimal measure of similarity in tomograms. This is particularly apparent for high binnings. Therefore, different, perhaps non-linear similarity metrics such as those used in machine learning can enhance template matching performance.^10,27,28,42^ However, for small macromolecules, generating adequate training data sets could be highly challenging, as manual curation would be limited by noise levels and visibility of macromolecules by eye. Second, analyzing lower-binned tomograms can potentially improve crosscorrelation-based template matching. The first developments have recently emerged in both 2D and 3D. 2D template matching^29,30^ avoids the high computational cost associated with exhaustive sampling, and, high resolution matching at low binning in 3D has recently been reported and demonstrated to give fewer false positive results.^35^ It also becomes evident that at such high binning, it is computationally most efficient to first use shape-based filtering, ideally with a spherical mask to filter candidate positions broadly^37^ and then further refine them locally with high-resolution template matching or filtering false positives by classification in programs such as RELION.^36^ To make shape-based picking easily accessible, we provide a Napari plugin via our software package pyTME^18^, which enables generating spherical, cylindrical, or ellipsoid templates and masks.

The high computational cost associated 3D template matching at low binning will be overcome in the future by further developing template matching software for efficient use on GPU without needing to bin the reconstructed tomograms^18,26^. Similarly higher angular sampling at lower binning might also be beneficial in specific cases^26,35^. Future developments will also need to tackle additional challenges such as noise, specimen motion resulting from problems with tilt alignment, sample deformation, and errors in CTF correction^29,43^.

Lastly, we suggest broadening benchmark entities beyond large, and highly abundant globular structures like the ribosome when evaluating new template-matching algorithms. Especially providing test sets of particles that have similar low-frequency information is necessary to determine the discriminatory power of novel templatematching methods, scores functions, or processing approaches. Novel methods should also be validated against the simple geometric shapes considered here to ensure they perform better and justify the higher computational cost.

## AUTHOR CONTRIBUTIONS

VM and MS conceived the study, performed research, wrote the initial draft, and edited the manuscript. JK provided feedback and edited the manuscript.

## V. DATA AVAILABILITY

The tomogram and ground truth picks are freely available at EMPIAR (EMPIAR-10988, TS_037). The scaled maps for the ribosome (EMD-3228), sphere, emoji and HA in various radii, the resulting picks, the raw data for plots, and used scripts are freely available on GitHub following link: https://github.com/maurerv/ribosomeTemplateMatching.

## ACKNOWLEDGMENTS

We thank Lukas Grunwald (Max Planck Institute for the Structure and Dynamics of Matter, Center for Free-Electron Laser Science (CFEL), Hamburg, Germany) for critical reading and editing of the theory section and Julia Mahamid (EMBL Heidelberg, Germany) for critical reading and feedback of the manuscript. We thank the EMBL IT and HPC resources for providing essential computational infrastructure. VM and JK acknowledge funding from the CSSB flagship project Plasmofraction. MS acknowledges support from a research fellowship from the EMBL Interdisciplinary Postdoc (EIPOD) Programme under Marie Curie Cofund Actions MSCA-COFUND-FP (grant agreement number: 847543).

## SUPPORTING INFORMATION

**FIG. S1.**
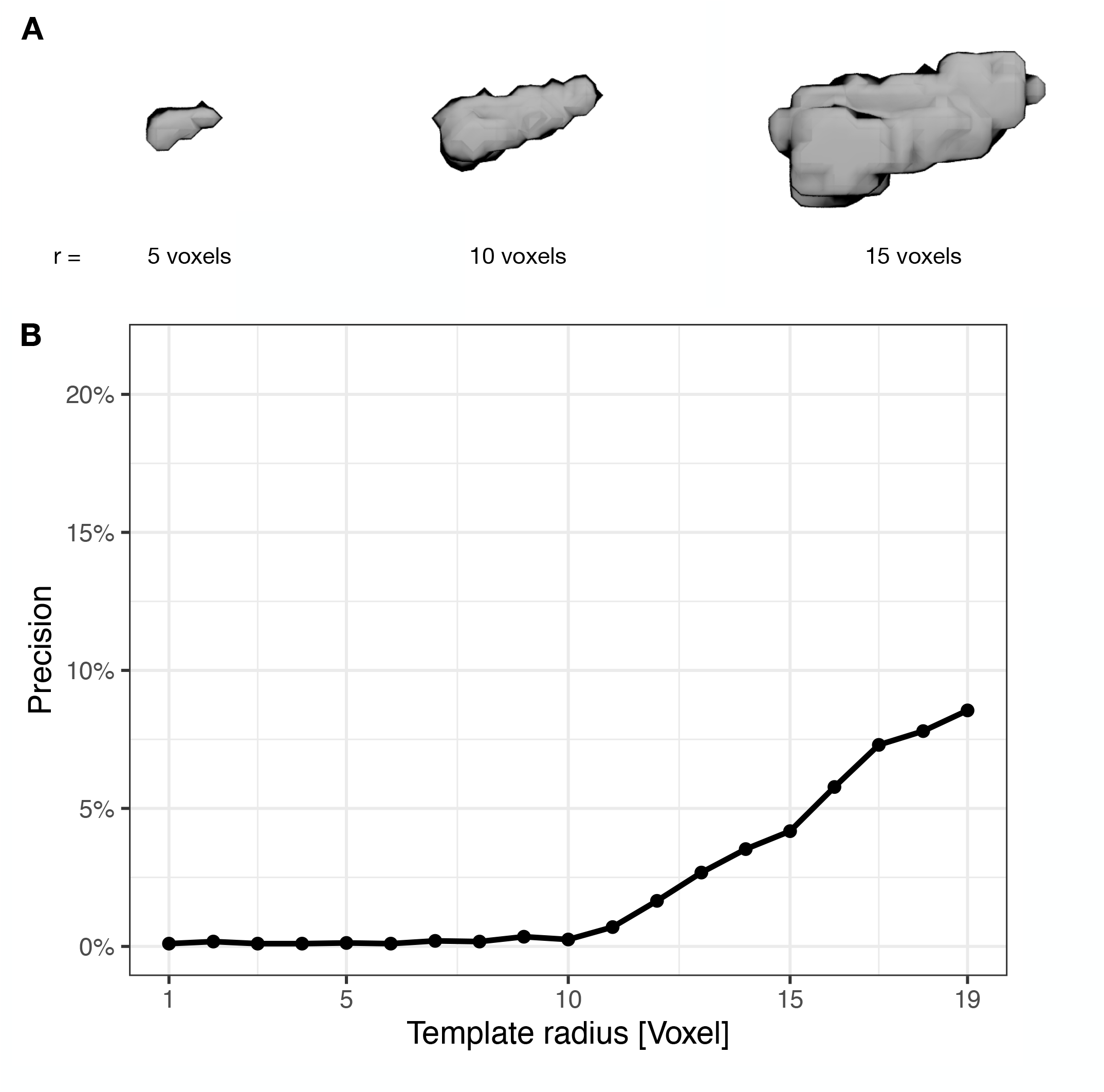
**A** Snapshots of the hemagglutinin (HA) templates with different radii used as a control. **B** Proportion of true positives out of all picked particles (precision) by the radius of templates. The HA structure was sampled with 1.944 angles.

